# The Proximal Centriole-Like Structure Anchors the Centriole to the Sperm Nucleus

**DOI:** 10.1101/2024.04.15.589606

**Authors:** Danielle B. Buglak, Kathleen H.M. Holmes, Brian J. Galletta, Nasser M. Rusan

## Abstract

Proper connection between the sperm head and tail is critical for sperm motility and fertilization. The link between the head and tail is mediated by the Head-Tail Coupling Apparatus (HTCA), which secures the axoneme (tail) to the nucleus (head). However, the molecular architecture of the HTCA is not well understood. Here, we use *Drosophila* to create a high-resolution map of proteins and structures at the HTCA throughout spermiogenesis. Using structured illumination microscopy, we demonstrate that key HTCA proteins Spag4 and Yuri form a ‘Centriole Cap’ that surrounds the centriole (or Basal Body) as it is inserted, or embedded into the surface of the nucleus. As development progresses, the centriole is laterally displaces to the side of the nucleus, during which time the HTCA expands under the nucleus, forming what we term the ‘Nuclear Shelf.’ We next show that the proximal centriole-like (PCL) structure is positioned under the Nuclear Shelf and functions as a critical stabilizer of the centriole-nuclear attachment. Together, our data indicate that the HTCA is complex, multi-point attachment site that simultaneously engages the PCL, the centriole, and the nucleus to ensure proper head-tail connection during late-stage spermiogenesis.

## INTRODUCTION

Fertilization requires a properly formed sperm, which is comprised of a ‘head’ containing the genetic material and a ‘tail’ that drives sperm motility. A stable connection between the sperm head and tail is critical for proper sperm motility and function. Failure to establish or maintain the connection between the head and tail can result in acephalic spermatozoa syndrome in patients, which presents with unstable connections between the sperm head and tail, including bent configurations and completely detached sperm heads, ultimately resulting in infertility (Baccetti et al., 1989, Chemes et al., 1987, Chemes et al., 1999, Perotti et al., 1981). Defects at the connection between the sperm head and tail have been linked to infertility across many other species, including bulls (Blom and Birch-Andersen, 1970), mice (Wang et al., 2022, Zhang et al., 2021), and *Drosophila* (Anderson et al., 2009, Kracklauer et al., 2010, Li et al., 2004, Sitaram et al., 2012, Texada et al., 2008).

The centriole, or Basal Body, is a critical structural component that links the sperm head and tail. The distal end of the centriole serves as a template for the axoneme that develops into the sperm tail, while the proximal end of the centriole helps anchor the tail to the nucleus (Galletta et al., 2020, Tates, 1971, Tokuyasu, 1975a). The connection between the head and tail at the centriole is known as the head-tail coupling apparatus (HTCA). Numerous proteins have been localized at or near the HTCA in various species, including nuclear envelope proteins such as the testis-specific SUN domain protein Spag4 and nuclear pore complexes (Tates, 1971, Kracklauer et al., 2010); centriolar proteins like CP110, CEP135, CETN1, FAM161A, and WDR90 (Subbiah, 2024; Khanal, 2021); and others including the gravitaxis protein Yuri Gagarin (Yuri) (Kracklauer et al., 2010, Texada et al., 2008), dynein and its adaptor Lis-1 (Li et al., 2004, Sitaram et al., 2012), the syntaxin interacting protein Salto (Augiere et al., 2019), the microtubule binding protein Hook1 (Kierszenbaum et al., 2011), the coiled-coil proteins CCDC42 and CCDC159 (Ge et al., 2024, Tapia Contreras and Hoyer-Fender, 2019), the sperm protein PMFBP1 (Zhu et al., 2018), and the linker protein CENTLEIN (Zhang et al., 2021). Mutations in many proteins localized to the HTCA are linked to head-tail connection defects and infertility. For example, mutations in the testis-specific SUN-domain protein SPAG4L (SUN5) and the sperm tail associated protein PMFBP1 account for nearly 70% of acephalic spermatozoa cases in humans (Zhang et al., 2021). In *Drosophila*, mutations in Spag4 and Yuri similarly lead to failed connection at the HTCA and infertility (Kracklauer et al., 2010, Texada et al., 2008). Although many different proteins have been localized to the HTCA and linked to infertility, it is unclear how these proteins assemble and function to physically link the sperm head and tail.

In *Drosophila*, sperm development begins with the asymmetric division of germline stem cells, giving rise to cells that undergo multiple divisions, eventually becoming spermatocytes (Demarco et al., 2014, Fuller, 1993) (**Figure S1A**). Spermatocytes then undergo meiosis to become spermatids that enter spermiogenesis, a developmental process that involves a series of dramatic morphological changes in which the axoneme elongates and the nucleus reshapes, ultimately resulting in mature sperm (**Figure S1A-C**) (Fabian and Brill, 2012, Tates, 1971, Tokuyasu, 1975a, Anderson, 1967, Shoup, 1967, Tokuyasu et al., 1972a, Tokuyasu et al., 1972b, Tokuyasu, 1974a, Tokuyasu, 1974b, Tokuyasu, 1975b, Tokuyasu et al., 1977, Stanley et al., 1972).

The connection between the nucleus and centriole is first established during early spermiogenesis and must be maintained as the spermatid develops (**Figure S1B-C**) (Fabian and Brill, 2012, Tates, 1971). Previous work discovered that initial attachment between the nucleus and centriole in Round spermatids is dependent on dynein at the nuclear envelope (Anderson et al., 2009, Li et al., 2004, Sitaram et al., 2012). In addition, restriction of pericentriolar material and microtubule nucleation to the proximal end of the centriole is necessary to ensure proper end-on attachment of the centriole to the nucleus (Galletta et al., 2020). However, less is known about how the nucleus and centriole maintain their connection as spermatids develop and remodel.

Here, we sought to create a high-resolution map of proteins and centriole structures present at the HTCA throughout spermiogenesis as the connection between the nucleus and centriole is established. Using structured illumination microscopy, we defined two novel structures at the HTCA during the remodeling that occurs during this period and identified a new role for the proximal centriole-like (PCL) structure, an atypical procentriole, in maintaining centriole positioning relative to the nucleus.

## RESULTS

### The HTCA is remodeled into a ‘Centriole Cap’ and ‘Nuclear Shelf’

While the HTCA is conceptually well understood, how it is established and develops in space and time during spermiogenesis is not well defined. Thus, the first goal was to perform structured illumination microscopy (SIM) on known HTCA proteins to define its structure throughout spermatid development. We selected the testis-specific SUN-domain protein Spag4 and gravitaxis protein Yuri, as both localize to the HTCA and mutations in each leads to sperm decapitation (Kracklauer et al., 2010, Texada et al., 2008), indicating a critical role in maintaining the connection between head and tail. In Early Round spermatids, the nucleus and centriole begin to form an attachment (**Figure S1B-C**). At this stage Spag4 encircled the entire nuclear envelope (**Figure 1A**). Shortly after, in Round spermatids, the centriole attached to the nucleus perpendicularly, or ‘end-on’ (**Figure S1B-C**), and Spag4 formed an asymmetrical crescent on one side of the nucleus, centered on the centriole (**Figure 1B**) (Kracklauer et al., 2010, Texada et al., 2008). Interestingly, Spag4 accumulated at higher levels in a small region on the nucleus at the centriole attachment point (**Figure 1B, arrow**). In Leaf stage spermatids, where the nucleus begins to elongate (**Figure S1B-C**), Spag4 dramatically remodeled into a structure tightly surrounding the proximal end of the centriole (**Figure 1C, F**), which begins to insert into an invagination of the nuclear surface (Tokuyasu, 1975a). We termed this structure the ‘Centriole Cap’. As the nucleus further elongated and narrowed in Late Leaf and Canoe stage spermatids (**Figure S1B-C**), the Centriole Cap lengthened (**Figure 1D-E, G**) in a manner corresponding with further insertion of the centriole into the nuclear invagination.

**Figure 1:**
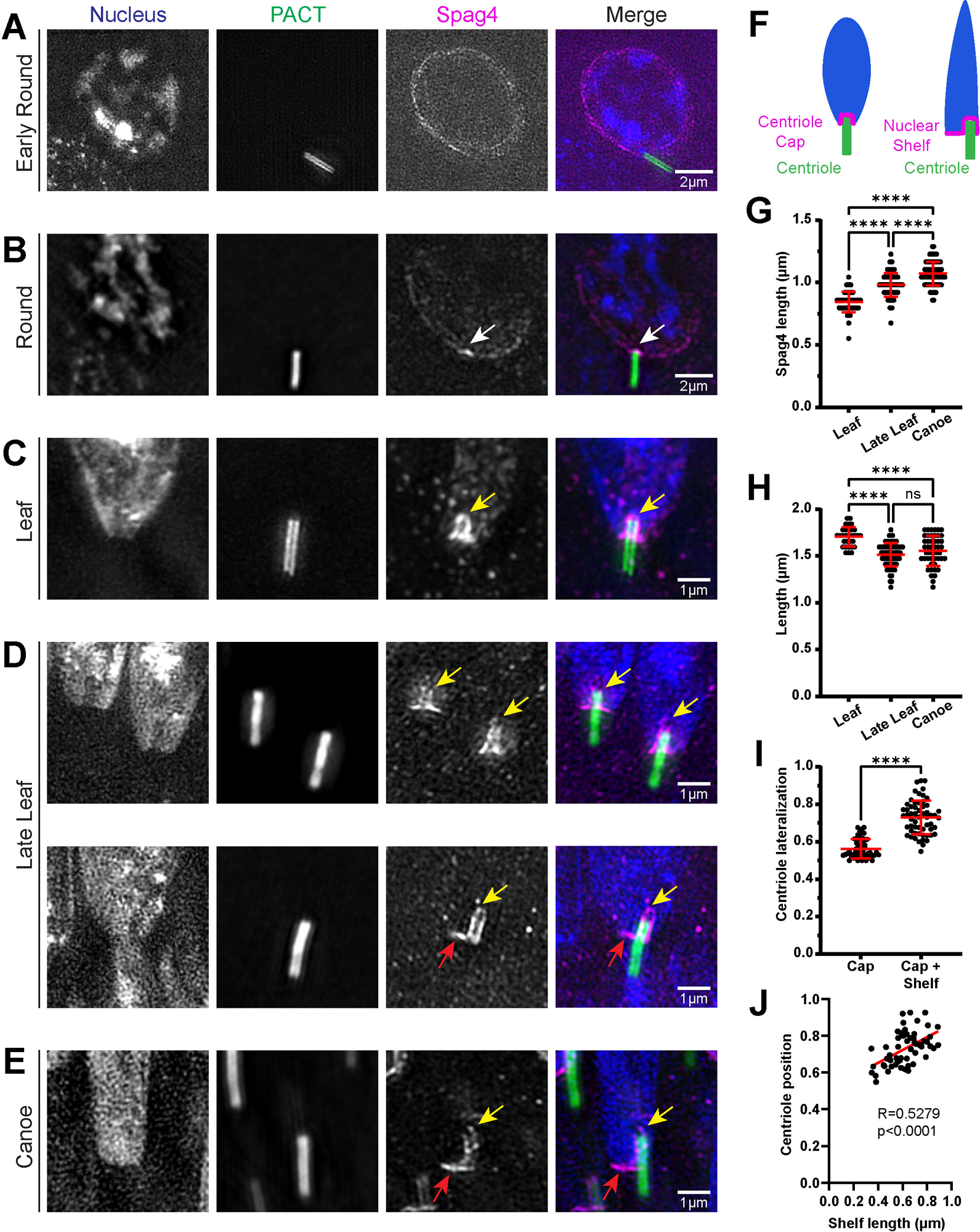
The HTCA is remodeled into a Centriole Cap and Nuclear Shelf. **(A-E)** Representative SIM images showing spermatids from wildtype testes during indicated developmental stages. Spermatids were labeled for the nucleus (DAPI, blue), centriole (PACT::GFP, green), and Spag4 (Spag4::6myc, magenta). White arrow denotes Spag4 accumulation at centriole docking site. Yellow arrows denote Centriole Caps. Red arrows denote nuclear shelves. Scale bars are labeled. **(F)** Cartoons depicting the Centriole Cap and Nuclear Shelf. **(G)** Quantification of Spag4 Centriole Cap length in Leaf (n=50), Late Leaf (n=78), and Canoe stage (n=85) spermatids. **(H)** Quantification of centriole length in Leaf (n=31), Late Leaf (n=72), and Canoe stage (n=47) spermatids. **(I)** Quantification of centriole lateralization index in spermatids with only a Centriole Cap (n=55) and spermatids with a Centriole Cap and Nuclear Shelf (n=58). **(J)** Scatter plot fit with a linear regression showing that centriole lateralization increases with Shelf length (n=56). ns=not significant, (****) p≤0.0001.

Measurements of centriole length during the centriole insertion process indicated that the centriole did not grow longer during nuclear insertion (**Figure 1H**). Thus the lengthening of the Centriole Cap is due to the centriole moving into the nuclear space rather than remaining fixed at a point and growing into the nucleus.

Following the Leaf and Late Leaf stages (**Figure 1C-D**), we observed nearly half of spermatids with asymmetrical localization of Spag4 by Canoe stage (**Figure 1E**). The asymmetry is a result of an extension of the HTCA to one side of the Cap and below the nucleus; we termed this structure the ‘Nuclear Shelf.’ The formation and positioning of the Shelf corresponded to the decentralization of the centriole relative to the nucleus (**Figure 1D-F**). Measuring the position of the centriole relative to the DNA signal along the short axis of the nucleus in Leaf through Canoe stage revealed that centrioles are positioned more laterally when a Shelf is present (**Figure 1I**). Furthermore, there is a positive correlation between Shelf length and the extent of centriole lateralization (**Figure 1J**), indicating that the events are coupled. Taken together, we show that the HTCA, as shown by Spag4 (**Figure 1**) and Yuri (**Figure S2**), forms an asymmetrical crescent that develops into a Centriole Cap surrounding the proximal end of the centriole as it inserts into the nucleus. The HTCA is then further remodeled by forming a Nuclear Shelf at the bottom of the nucleus while the centriole is laterally positioned to one side of the nucleus.

### The centriole adjunct is extensively remodeled as the Centriole Cap elongates

Given the extensive remodeling of the HTCA, we sought to investigate other structures associated with spermatid centrioles during spermiogenesis. One such structure is the centriole adjunct (CA), a dense structure of pericentriolar material that surrounds the centriole and nucleates a specialized microtubule network called the manchette, which is important for spermatid nuclear shaping (Riparbelli et al., 2020, Russell et al., 1991). Electron micrographs show that the CA is positioned near the site of the HTCA in developing spermatids (Riparbelli et al., 2020, Tokuyasu, 1975a); thus, we hypothesized that the CA associates with, and possibly regulates Centriole Cap or Nuclear Shelf formation. To investigate CA dynamics, we immunostained spermatids for Asterless (Asl), a known regulator of the PCM and centriole length, and component of the CA (Varmark, 2007; Galletta et al., 2016, Blachon et al., 2008). As previously shown, we found that Asl localized along the entire length of the centriole in Round spermatids (**Figure 2A**) (Galletta et al., 2016, Khire et al., 2015). Asl then began to condense into a ‘collar’ and then a ‘ring’ in Leaf and Canoe stage spermatids, respectively (**Figure 2B-C**). The transition of the CA to the collar and ring perfectly correlated with the insertion of the centriole into the nucleus (**Figure 2B-D**) and formation and elongation of the Centriole Cap shown by Spag4 (**Figure 2E-F**). Finally, Asl, was further remodeled asymmetrically in late Canoe spermatids to one side of the centriole, positioned beneath the Spag4 Nuclear Shelf (**Figure 2D**). Thus, we hypothesize that condensation of the CA into a ring exposes the proximal end of the centriole and facilitates centriole insertion into the nucleus and remodeling of the HTCA. Unfortunately, we were unable to directly test whether the CA is important for insertion or important for HTCA formation as *asl* mutants also affect centriole duplication and microtubule nucleation (Blachon, 2008; Varmark, 2007).

**Figure 2:**
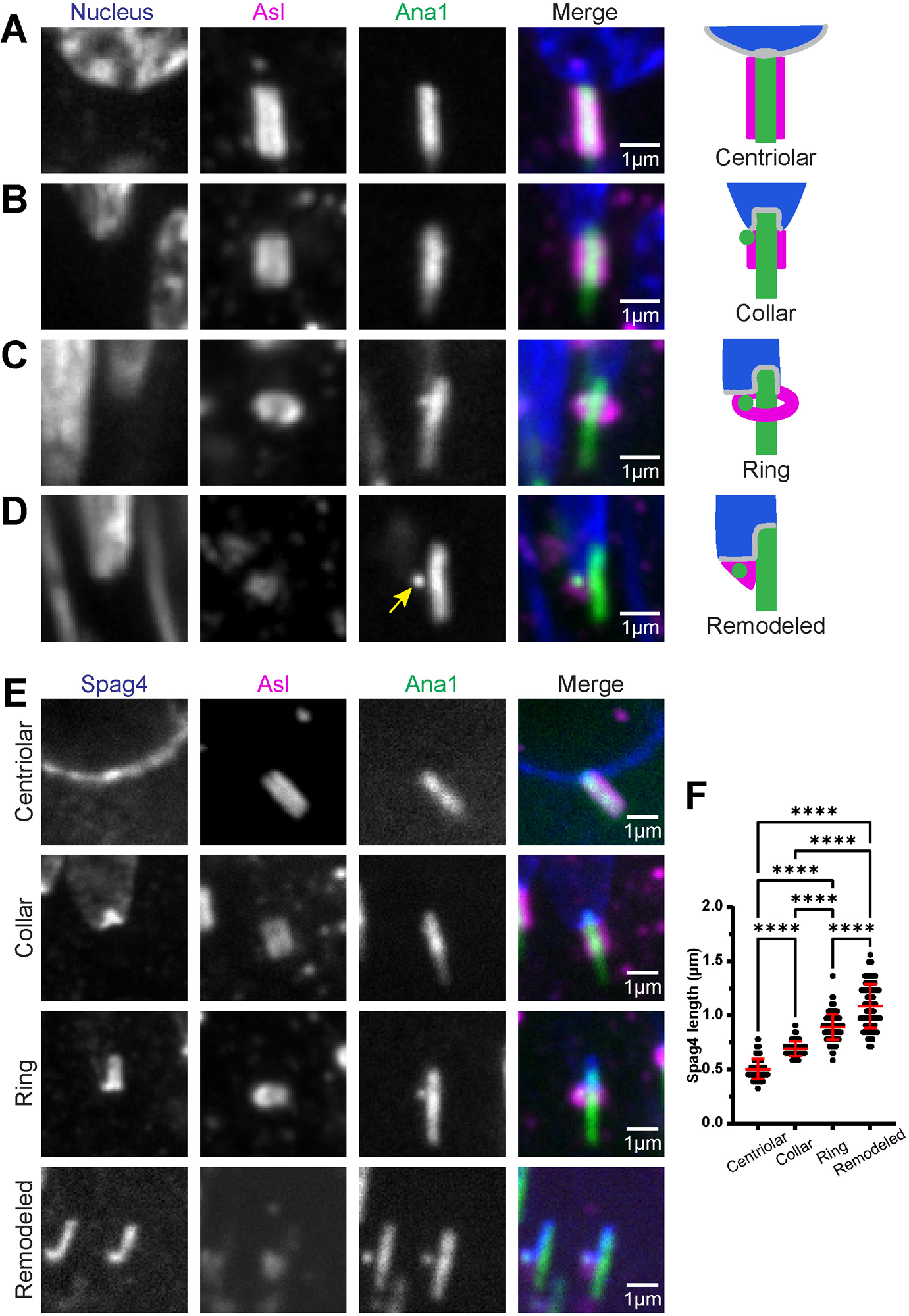
The Centriole Adjunct is extensively remodeled as the Centriole Cap elongates. **(A-D)** Representative images showing spermatid centrioles from wildtype testes at indicated stages of CA remodeling (right). Spermatids were labeled for the CA (Asl, magenta) and centriole/PCL (Ana1::tdTomato, green). Yellow arrow denotes PCL. Scale bar, 1μm. Cartoons represent different stages of CA remodeling. **(E)** Representative images showing spermatids from wildtype testes at indicated stages of CA remodeling. Spermatids were labeled for Spag4 (Spag4::6myc, green), centriole/PCL (Ana::tdTomato, magenta), and CA (Asl, gray). Scale bar, 1μm. **(F)** Quantification of Spag4 Centriole Cap length in spermatids at centriolar (n=155), collar (n=147), ring (n=134), and remodeled (n=110) stages of CA remodeling. (****) p≤0.0001.

### The PCL is associated with the Nuclear Shelf

As the centriole is positioned laterally on one side of the nucleus, the CA and the Centriole Cap become asymmetrical (**Figure 1D, 2D**). Further analysis of the CA in late Canoe states revealed an Ana1 (centriole marker) positive structure within the CA (**Figure 2D**). This structure is known as the proximal centriole-like (PCL) structure, an atypical procentriole found in spermatids that serves as a template for daughter centriole formation during the first zygotic divisions (Blachon et al., 2009, Blachon et al., 2014). Furthermore, mutations in PCL-associated proteins lead to male infertility, immotile sperm, and sperm structural abnormalities (Khire et al., 2016). This led us to hypothesize that the PCL is important in defining the Nuclear Shelf and creating a stable HTCA. To test this hypothesis, we first determined the position of the PCL relative to the nucleus, centriole, Centriole Cap, and Nuclear Shelf. We utilized transgenic flies expressing Sas-6::TagRFP, which labels both the proximal end of the centriole and the PCL (Galletta et al., 2016, Jo et al., 2019). We found that the position of the PCL was nearly indistinguishable from the proximal end of the centriole in Leaf stage spermatids, but became displaced from the proximal end as spermatids transitioned into Late Leaf and Canoe stages (**Figure 3A**). We measured the distance between the proximal end of the centriole and the PCL at each stage and found that the Sas6-positive PCL moves further away from the base of the centriole as it is inserted into the nucleus (**Figure 3B**).Thus, throughout the centriole insertion process, the PCL was always positioned at the base of the nucleus, eventually settling into a location below the Nuclear Shelf in Canoe stage spermatids (**Figure 3C**) and subsequently at the base of the needle shaped nuclei in mature sperm (**Figure S1B**).

**Figure 3:**
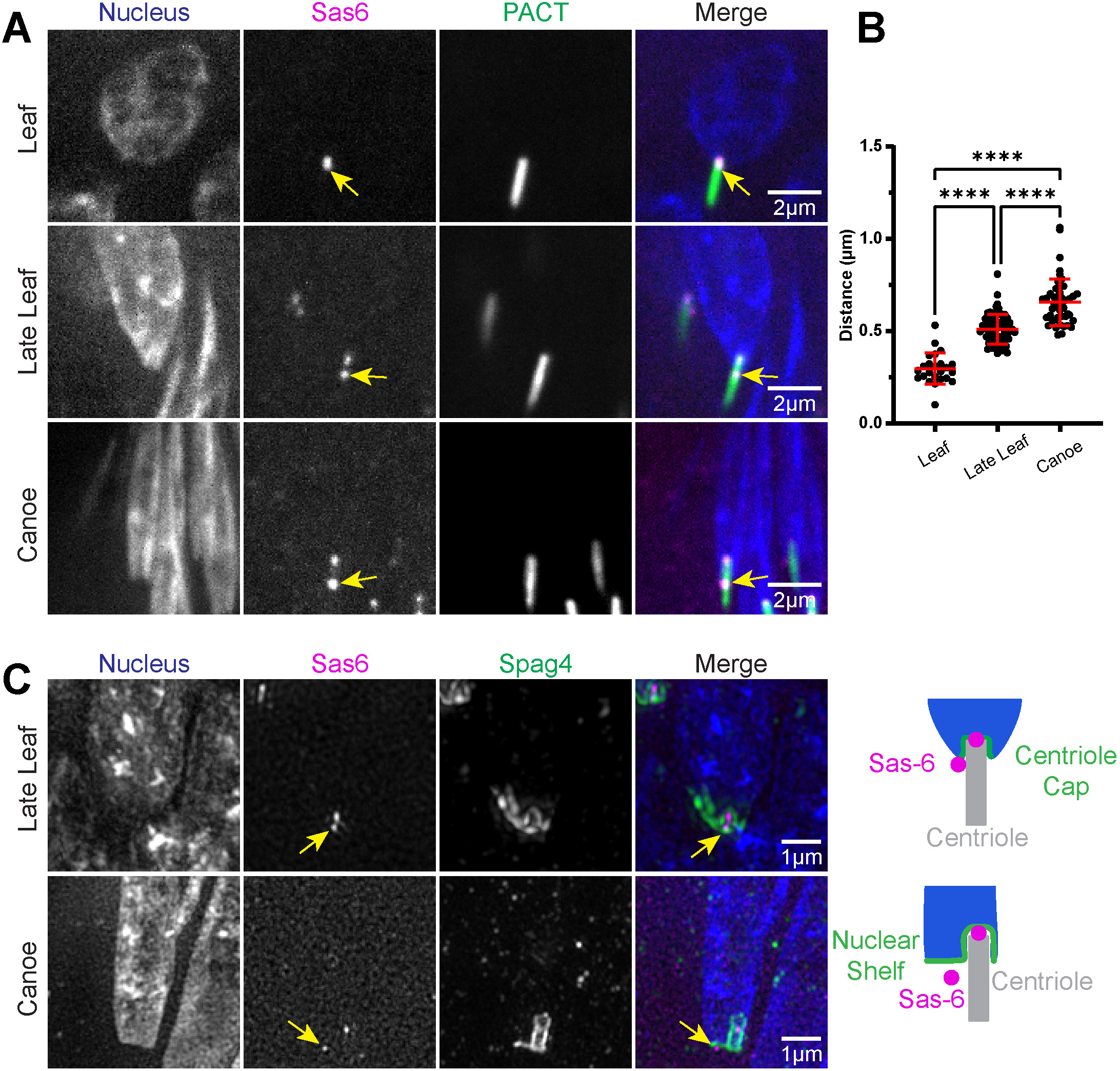
The PCL is associated with the Nuclear Shelf during centriole insertion. **(A)** Representative images showing spermatids from wildtype testes during indicated developmental stages. Spermatids were labeled with the nucleus (DAPI, blue), centriole (PACT::GFP, green), and Sas6 marking the proximal end of the centriole and the PCL (TagRFP::Sas6, magenta). Scale bar, 2μm. Yellow arrows denote PCL. **(B)** Quantification of the distance between the proximal end of the centriole and the PCL in Leaf (n=24), Late Leaf (n=67), and Canoe (n=46) stage spermatids. **(C)** Representative SIM images showing spermatids from wildtype testes during indicated developmental stages. Spermatids were labeled with the nucleus (DAPI, blue), Spag4 (Spag4::6myc, green), and Sas6 marking the proximal end of the centriole and the PCL (TagRFP::Sas6, magenta). Yellow arrows denote PCL. Scale bar, 1μm. Cartoons depict position of the PCL beneath the Centriole Cap and Nuclear Shelf. (****) p≤0.0001.

Taken together, the PCL is initially formed at the base of the centriole, then it remains associated with the remodeled CA collar and ring in a position below the nucleus where it ultimately becomes a part of the final mature sperm structure.

### The PCL acts as an anchor during centriole insertion

Given that both the CA and PCL are positioned below the Nuclear Shelf and remodeled during centriole insertion, we hypothesized that the CA or the PCL is necessary for proper assembly or maintenance of the HTCA. Unfortunately, we were unable to directly test the CA, but we were able to genetically eliminate the PCL using a mutant for the centriole protein Poc1. The *Poc1* gene encodes two protein isoforms, Poc1A, a centriole protein important for centriole elongation, and Poc1B, a centriolar protein that is highly expressed in the testes that exclusively localizes to the PCL (Khire et al., 2016; **Figure S3**).

To test the role of the PCL in sperm head/tail connection, we analyzed *poc1^W87X^* mutant flies previously shown to disrupt the architecture of the PCL (Khire et al., 2016). In Round spermatids, the nucleus and centriole joined to establish the HTCA in both wildtype and *poc1* mutants (**Figure 4A; Figures S4A**). Furthermore, in both mutants and controls, the CA was properly localized along the entire centriole length, and Spag4 still formed both a crescent with the expected accumulation at the site of centriole docking (**Figure 4A; Figure S4A**).In addition, no significant difference in Spag4 cap length was detected in *poc1* mutants through Canoe stage spermatids that contained collar and ring shaped CAs (**Figure 4B,C,E; Figure S3B,C**). *poc1* mutants also properly formed Spag4 Nuclear Shelves and laterally positioned centrioles (**Figure 4C; Figure S4C**), indicating that centriole lateralization and asymmetry of the HTCA is not dependent on the PCL. However, during CA remodeling from a ring to a density beneath the Nuclear Shelf (**Figure 2D**), *poc1* mutant spermatids displayed shorter Spag4 Centriole Caps (**Figure 4D-E; Figure S4D**). We also tested a hypomorph *poc1* allele (*poc1^c06059^*) that reduces Poc1 expression (Khire et al., 2016), and similarly found shorter Spag4 Cap length in late stage spermatids (**Figure S5A-B**). We therefore conclude that the PCL is not required for centriole insertion and lateralization, but is required for maintaining insertion of the centriole in late stages of spermiogenesis.

**Figure 4:**
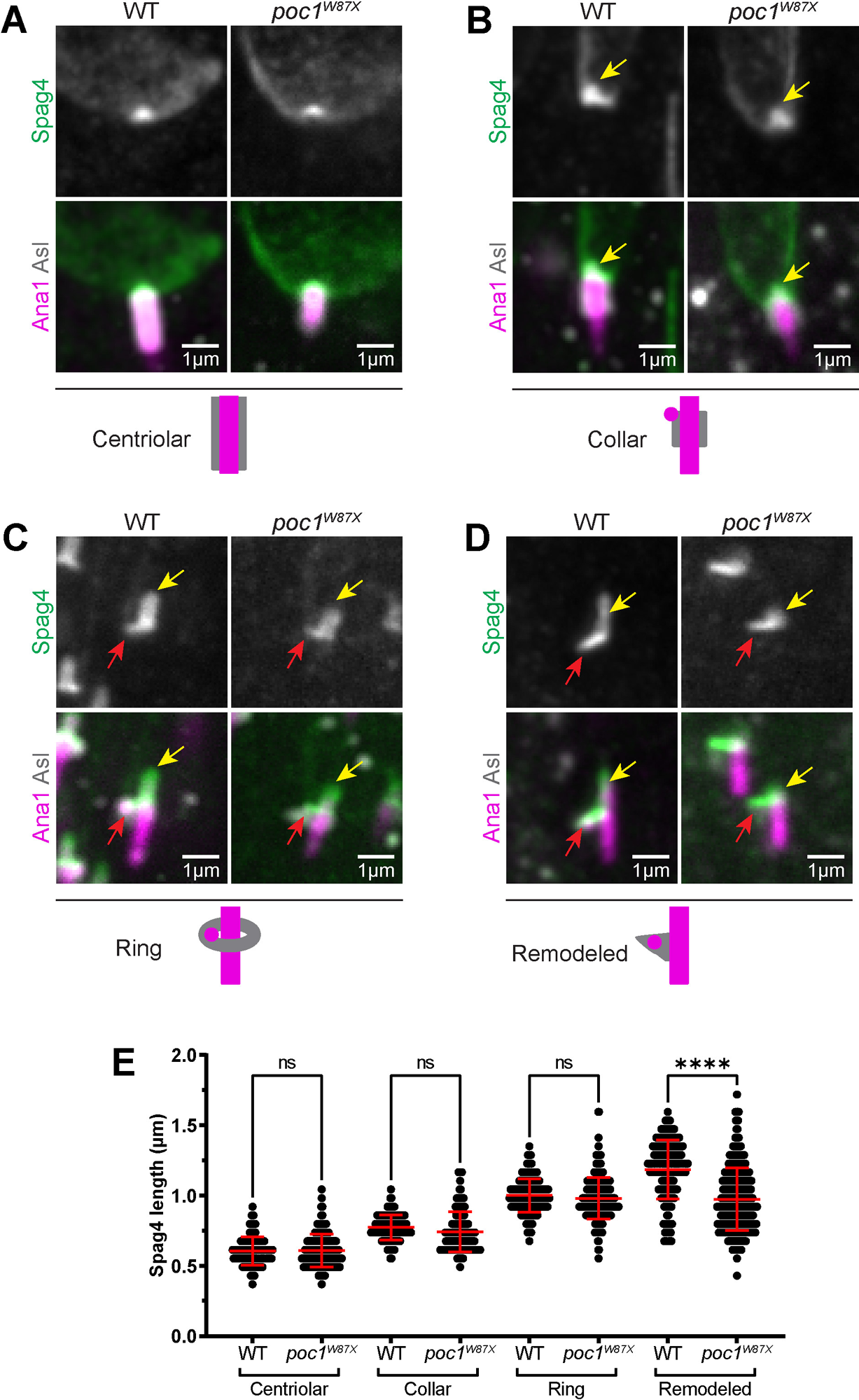
*poc1* mutants have shorter Centriole Caps following remodeling of the CA. **(A-D)** Representative images showing spermatids from wildtype (left) and *poc1^W87X^*mutant (right) testes at indicated stages of CA remodeling. Spermatids were labeled with Spag4 (Spag4::6myc, green), centriole/PCL (Ana1::tdTomato), and the CA (Asl, gray). Yellow arrows denote Centriole Cap. Red arrows denote Nuclear Shelf. Cartoons depict different stages of CA remodeling. Scale bar, 1μm. **(E)** Quantification of Centriole Cap length in wildtype and *poc1^W87X^*mutant spermatids during centriolar (wildtype n=86; mutant n=148), collar (wildtype n=125; mutant n=142), ring (wildtype n=258; mutant n=261), and remodeled (wildtype n=188; mutant n=322) stages of CA remodeling. ns=not significant, (****) p≤0.0001.

To ensure that the loss of centriole insertion was due to disruption of the PCL and not due to other effects of *poc1* manipulation, we also disrupted the PCL via *plk4* RNAi, as formation of the PCL is dependent on Plk4 (Blachon et al., 2009). Knockdown of Plk4 did not entirely eliminate the PCL in all spermatid as evidenced by the presence of Sas-6 signal at the PCL (**Figure S5E**). However, we found that spermatids with centrioles that were not properly inserted or were completely detached had significantly reduced Sas-6 at the PCL, relative to Sas-6 at the proximal end of the centriole, in contrast to controls (and compared to *plk4* RNAi spermatids with normal centriole insertion) (**Figure S5E-F**). Together, our data is consistent with a model whereby Poc1 and the PCL are not necessary for centriole lateralization and establishment of the Nuclear Shelf, but are critical for maintaining centriole insertion and anchoring the centriole during later stages.

Given that *poc1* mutant spermatids had significantly shorter Spag4 Caps in later stages of development, we hypothesized that centrioles were no longer stably linked to the nucleus. In wildtype spermatids, the majority of centrioles were laterally and perpendicularly attached to the nucleus (**Figure 5A,B**). However, only ∼50% of *poc1* mutant spermatids were normally inserted into the nucleus (**Figure 5A,B**). In the other 50% of cases, *poc1* mutant centrioles were not properly inserted, attached at abnormal angles (bent), or completely detached from the nucleus (**Figure 5A-B**). *poc1^c06059^*flies similarly had reduced centriole insertion compared to controls, though to a lesser extent than *poc1^W87X^* flies (**Figure S5C,D**).

**Figure 5:**
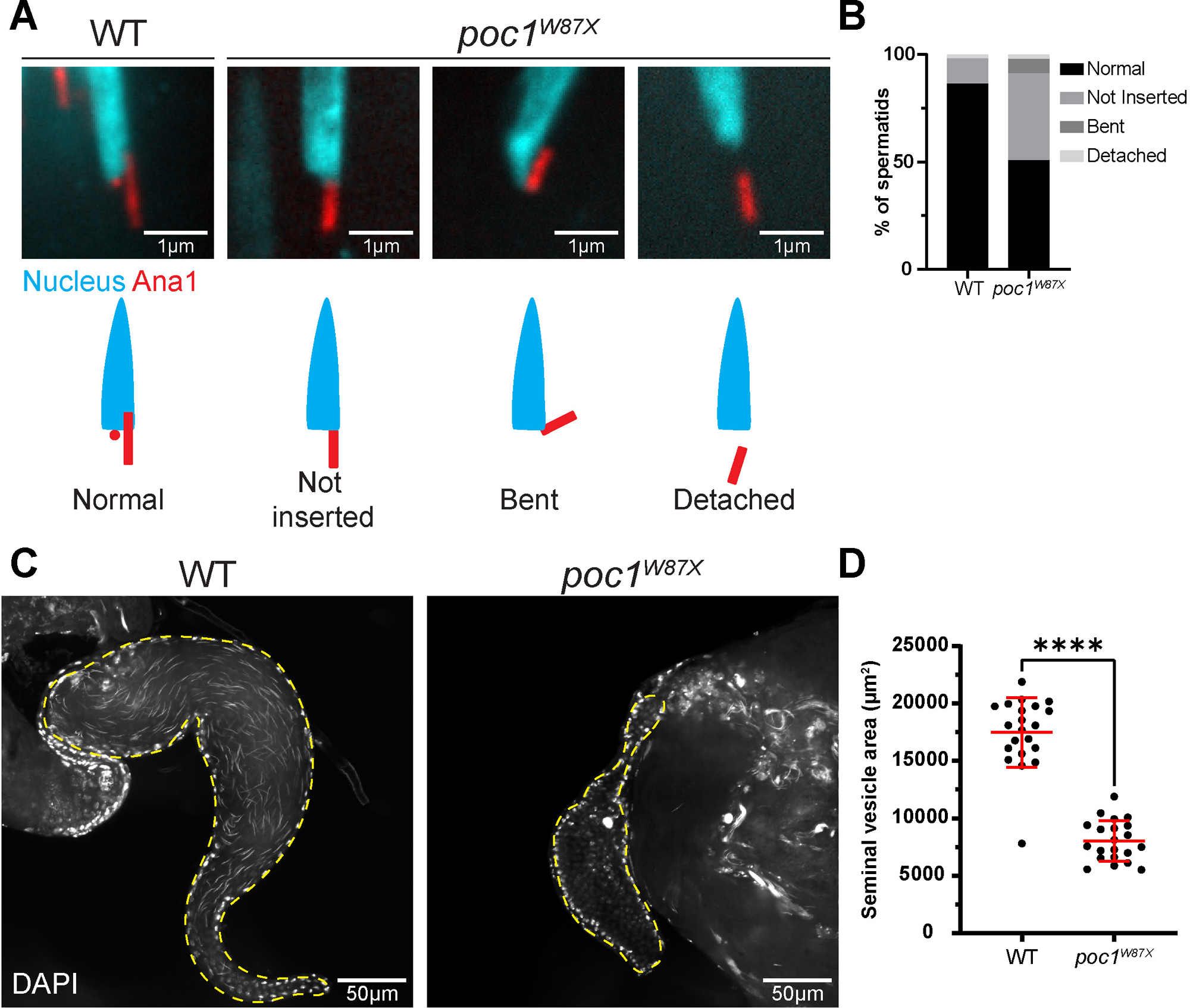
The PCL is required to maintain centriole insertion in late stages of spermiogenesis. **(A)** Representative images showing wildtype (left) and *poc1^W87X^*mutant (right) Canoe stage spermatids with various HTCA phenotypes. Spermatids were labeled with the nucleus (DAPI, cyan) and the centriole/PCL (Ana1::tdTomato, red). Scale bar, 1μm. Cartoons depict centrioles that are normal, not inserted, bent, or detached relative to the nucleus. **(B)** Quantification of wildtype (n=367) and *poc1^W87X^* mutant (n=597) spermatids with various HTCA phenotypes. **(C)** Representative images showing wildtype (left) and *poc1^W87X^* mutant (right) seminal vesicles. Mature sperm were labeled with the nucleus (DAPI). Scale bar, 50μm. **(D)** Quantification of seminal vesicle area in wildtype (n=21) and *poc1^W87X^* mutant (n=21) testes. (****) p≤0.0001.

These data suggest that loss of the PCL results in a stepwise deterioration of the centriole-nuclear attachment, starting with a regression from the nucleus, followed by a weakening of the connection, and finally full centriole detachment. We thus hypothesized that without proper attachment the sperm’s ability to swim and reach the seminal vesicle would be reduced. Indeed, explants of seminal vesicles revealed that *poc1* mutants had significantly smaller seminal vesicles with little to no sperm compared to controls (**Figure 5C-D**). This is fully consistent with the previous observation that flies with mutations in *poc1* lead to immotile sperm and sterility (Khire et al., 2016).

## DISCUSSION

Mature sperm must form a stable connection between the head and tail. This linkage is mediated by the HTCA, which connects the sperm nucleus to the axoneme via the centriole. In *Drosophila*, initial attachment of the centriole is end-on to the nucleus, while the final mature sperm form stable lateral attachment. During the centriole transition from end-on to lateral attachment, the HTCA is remodeled while maintaining the connection between nucleus and the centriole. Our study used structured illumination microscopy to characterize remodeling of the HTCA and HTCA-associated structures during spermiogenesis. We have discovered a new role for the PCL as a structural anchor that maintains centriole positioning and stable sperm tail attachment.

Our data, combined with previous studies, are consistent with a four stage process of HTCA development (**Figure 6**), divided into two major phases. First is the ‘Establishment’ phase, which consists of nuclear ‘Search’ by the centriole (stage 1) followed by centriole ‘Attachment’ to the nucleus (stage 2). This establishment phase is mediated by dynein and its adaptors Lis-1 and Asunder, and requires restriction of the PCM and microtubules to the proximal end of the centriole to ensure end-on attachment (Anderson et al., 2009, Galletta et al., 2020, Li et al., 2004, Sitaram et al., 2012). Importantly during this time, the CA, as marked by Asl, is positioned along the entire length of the centriole (**Figure 6, Pink**). Next is the ‘Maintenance’ phase, which consists of centriole ‘Insertion’ into the nucleus (stage 3) followed by centriole lateralization (stage 4). During centriole insertion, the CA begins to condense and expose the proximal end of the centriole as Spag4 and Yuri form a ‘Centriole Cap’ around the centriole as it is inserted. As the centriole moves laterally (still remaining perpendicular to the nucleus), Spag4 and Yuri localization expands to the ‘Nuclear Shelf’ along the bottom of the nucleus.

**Figure 6:**
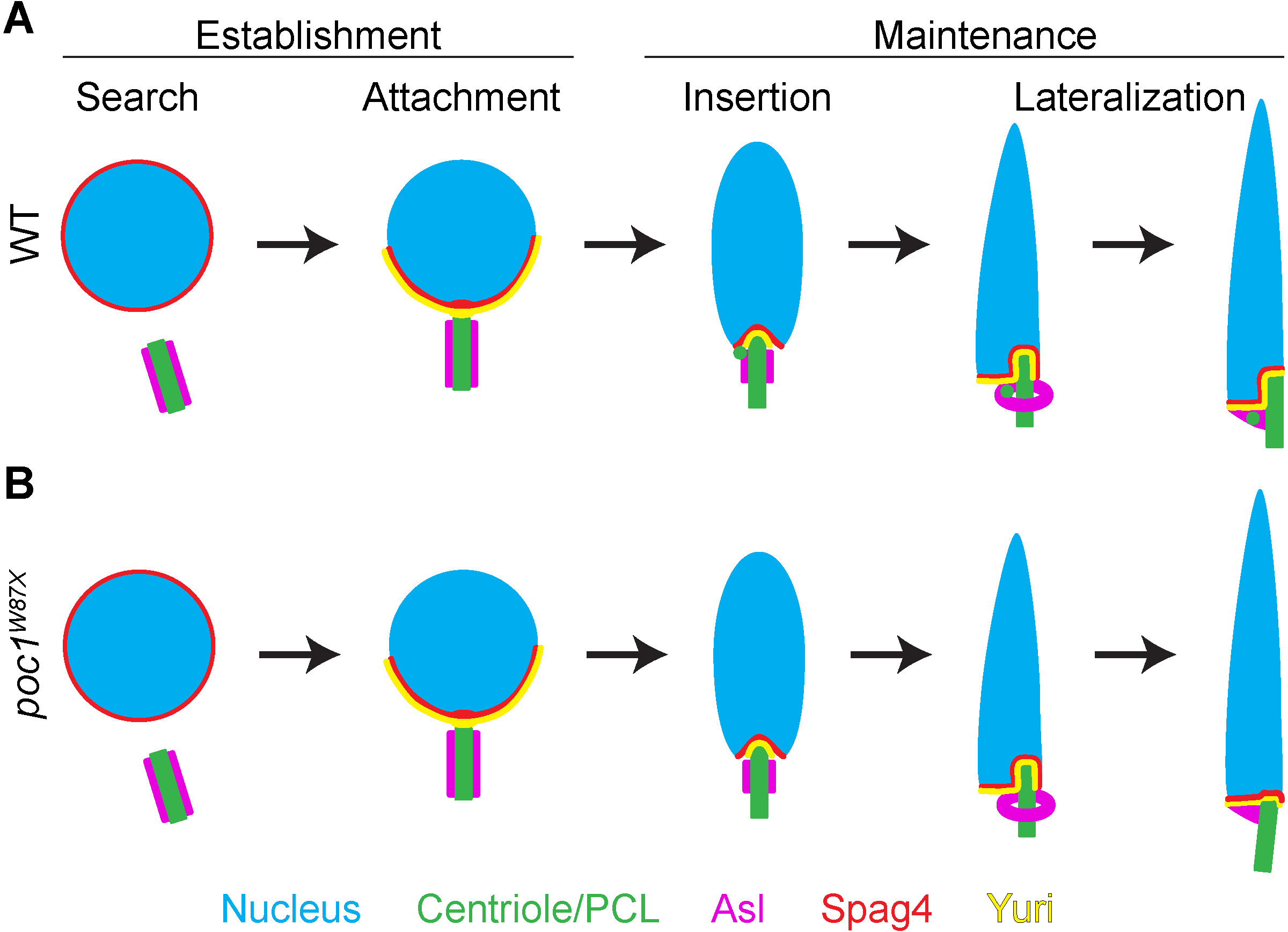
Model of HTCA establishment and remodeling. **(A)** In normal development, the nucleus and centriole use microtubules and dynein (not depicted) during the Search phase to initiate their interaction, which leads to the direct centriole-nucleus interaction during the Attachment phase. This constitutes Establishment of the HTCA. During Attachment, spermatid nuclei are Round, the CA is localized along the length of the centriole, and Spag4 and Yuri form a crescent with accumulation at the site of centriole docking. Development proceeds with the Maintenance phase during which the centriole undergoes Insertion into the nucleus at Leaf stage spermatids. Spag4 and Yuri form a Centriole Cap surrounding the inserting centriole and the CA begins to condense into a collar. As Insertion continues into Canoe stage, the centriole undergoes Lateralization during which the centriole moves to one side of the nucleus. On the opposite side, Spag4 and Yuri form a Nuclear Shelf, with the PCL localized beneath the Shelf. The CA further condenses into a ring as the centriole continues to insert into the nucleus. In the final stages of Lateralization, the CA condenses around the PCL to sit beneath the Nuclear Shelf. This results in a stable lateral connection between the sperm head and tail. **(B)** In *poc1* mutants, Search, Attachment, and Insertion all occur, with normal remodeling of the CA and elongation of the nucleus. However, in the final stages of Lateralization, centrioles are not anchored by the PCL and lose their insertion, resulting in an unstable attachment between head and tail.

Concomitant with insertion and lateralization of the centriole is the formation and movement of the PCL, and restructuring of the CA. Presumably, these events are coordinated to ensure that the final position of the CA and PCL is below the Nuclear Shelf. We hypothesized that the PCL would be critical for insertion and/or lateralization of the centriole in the nucleus centriole positioning relative to the nucleus during HTCA formation. We therefore investigated *poc1* mutants, which have abnormal PCL structures. Interestingly. spermatids in these mutants are able to progress through centriole insertion and lateralization, indicating that the PCL does not play a role in these early steps of HTCA formation. However, these spermatids are unable to maintain centriole insertion into the nucleus in late stages of spermiogenesis, indicating a role for the PCL in stabilizing the final head-tail connection. We propose that the HTCA forms a multi-point, L-shaped attachment surface that links the centriole, PCL and the nucleus. Such an expanded attachment site might be critical for resisting forces of the swimming tail and ensuring its stable connection to the head. This is consistent with our finding that *poc1* mutant flies have little to no sperm in their seminal vesicles, as sperm must be able to swim in order to reach the seminal vesicle. In humans, sperm with abnormal morphology have reduced expression of *POC1B* mRNA (Platts et al., 2007), and abnormal levels of POC1B at the centrioles were found in patients with unexplained infertility (Turner et al., 2021). Furthermore, the atypical centriole in human and bovine sperm has an enrichment of POC1B {Fishman, 2018 #53} (Khanal et al., 2021). Interestingly, at least in bovine sperm, this atypical centriole appears to be important creating a specialized coupling between the head and tail to control movement (Khanal et al., 2021). Together these observations all for the hypothesis that atypical centrioles like the PCL are necessary for proper sperm motility and fertility across species.

One interesting aspect of our study was the relationship between the CA and the nucleus. Several previous studying have shown that the CA at some point forms a ring structure (Blachon, 2008; Galletta, 2016; Tates, 1971), however not much more was known. By careful timing and correlation with centriole dynamics, we observed that the CA localized along the entire centriole through the centriole ‘Attachment’ phase. Then, the CA began to remodel by moving toward the longitudinal center of the centriole, exposing both the distal and proximal ends. The mechanism by which the CA condenses into a focused ring and then a density under the nucleus is completely unknown. However, the CA remodeling could involve a phase transition centered around the PCL as it moves from its initial position at the distal end fothe centriole to its final position in the middle of the centriole, just below the nuclear shelf. Future work to test the role of the PCL in forming the CA will require mutations that would completely eliminate the PCL rather than disrupt its structure as seen in *poc1* and *sas6* mutations. If such a mutant can be found, it will allow one to prevent CA remodeling and prevent the exposure of the proximal end of the centriole, which will probe the relationship between centriole exposure and insertion. In addition, mutations that disrupt CA formation that do not affect other aspects of spermiogenesis will be important to identify. Such CA mutations will allow one to test several hypotheses such as the possible role of CA condensation upstream of PCL movement along the centriole, or to investigate if PCL movement without the CA is sufficient for driving proper centriole insertion.

Spag4 and Yuri have been shown to be critical for the HTCA as mutations in either lead to centriole detachment from the nucleus and infertility (Kracklauer et al., 2010, Texada et al., 2008). This leads to the hypothesis that Spag4 and Yuri function in the same pathway. In support of this, we found that they have identical localizations and dynamics during HTCA remodeling – they both form a Centriole Cap and Nuclear Shelf. However, it is unclear how Spag4 and Yuri function, independently or in unison. Future work will focus on their precise function and the identification of proteins that might link the inner-nuclear membrane protein Spag4 with the cytoplasmic protein Yuri. More broadly, a more complete understanding of the molecular composition of the HTCA will be required to fully investigate HTCA formation and function in the critical head-tail connection process.

## ACKNOWLEDGEMENTS

We thank Xufeng Wu and the NHLBI Light Microscopy Core for assistance with SIM imaging; Carey Fagerstrom for *Drosophila* cloning; Tomer Avidor-Reiss and the Bloomington *Drosophila* stock center for flies; Kathleen Beckingham for antibodies; Alex Kelly, Takashi Akera, Leah Rosin, Matthew Hannaford, Chaitali Khan, Ryan O’Neill, and Solo Aviles for helpful discussion. This work is supported by the Division of Intramural Research at the NHLBI/NIH (1ZIAHL006126 to NMR).

## METHODS

### Flies

*D. melanogaster* were maintained and crosses were performed at 25°C. UAS-Ana1::tdTomato expresses Ana1 at endogenous levels without the use of Gal4 (Blachon et al., 2008). We used the following strains: *y,w*, *UAS-Ana1::tdTomato, ubi-PACT::GFP* (Galletta et al., 2020, Martinez-Campos et al., 2004)*, Spag4::6myc* (Bloomington #29981, #29980), *TagRFP::Sas-6* (Galletta, 2020), *poc1A::GFP* (T. Avidor-Reiss), *poc1B::GFP* (T. Avidor-Reiss), *poc1^W87X^* (T. Avidor-Reiss)*, poc1^c06059^*(T. Avidor-Reiss), and *plk4* RNAi (Bloomington #57221).

### Testis fixation and immunofluorescence

Testes from 1-3 day old males were dissected in Schneider’s media with antibiotic-antimycotic and fixed in 9% paraformaldehyde (PFA) at room temperature (RT) for 15-20 min. Testes were washed in PBS with 0.3% Triton X-100 (PBST) then blocked for 2-6 hours at RT in PBST with 5% normal goat serum. Samples were incubated in primary antibody in blocking solution at 4°C overnight, washed 3 times for 10 min in PBST, then left in secondary antibody in blocking solution for 6-8 hours at RT. Following three 10 min washes in PBST, samples were mounted in AquaPolymount for confocal imaging or Vectashield for SIM. For visualization of Yuri, pharate adults were dissected as above and fixed in 3% PFA at RT for 10 min. Staining was then performed as above.

| Antibody/Stain | Type | Source | Catalog # | Dilution |
| --- | --- | --- | --- | --- |
| Mouse anti-9E-10 c-myc | Primary | DHSB | 9E 10-c | 1:1,000 |
| Guinea pig anti-Asl | Primary | (Klebba et al., 2013) |  | 1:10,000 |
| Chicken anti-Yuri 4012 | Primary | (Texada et al., 2008) |  | 1:100 |
| AlexaFluor488 conjugated | Secondary | Thermo Fisher Scientific |  | 1:1,000 |
| AlexaFluor568 conjugated | Secondary | Thermo Fisher Scientific |  | 1:1,000 |
| AlexaFluor647 conjugated | Secondary | Thermo Fisher Scientific |  | 1:1,000 |
| DAPI | N/A | Thermo Fisher Scientific | D1306 | 1:100 –<br>1:1,000 |
| Hoescht 33342 | N/A | Life Technologies | H3570 | 1:100 |

### Confocal imaging

Confocal images were primarily acquired using a Nikon Eclipse Ti2 (Nikon Instruments) with a Yokogawa CSU-W1 spinning disk confocal head (Yokogawa, Life Science) equipped with a Prime BSI cMOS camera (Teledyne Photometrics) and Nikon Elements software (Nikon Instruments). Spermatids were imaged using a 100x/1.35 NA silicone immersion oil objective or 100xTIRF/1.49 NA oil immersion objective. Some images were acquired using a Nikon Eclipse Ti2 (Nikon Instruments) with a CSU-22 spinning disk confocal head (Visitech International) equipped with an ORCA-Flash 4.0 CMOS camera (Hamamatsu Photonics) and MetaMorph software (Molecular Devices) or a Nikon Eclipse Ti2 (Nikon Instruments) with a Yokogawa CSU-X1 spinning disk confocal head (Yokogawa, Life Science) equipped with an OCRA-Flash 4.0 CMOS camera (Hamamatsu Photonics) and Nikon Elements software (Nikon Instruments). Spermatids were imaged using a 100xTIRF/1.49 NA oil immersion objective; seminal vesicles were imaged using a 40x/1.30 NA oil immersion objective. 405, 488, 561, and 641nm laser lines were used on all systems. All images were analyzed and processed in FIJI (ImageJ, National Institutes of Health).

### Structured illumination microscopy

SIM images were acquired using an OMX4 (GE Healthcare) with DeltaVision Immersion Oil 1.514 and a **60x 1.42 NA oil immersion objective** (Olympus). To ensure successful reconstruction, laser settings for each channel were adjusted to produce a 1000 unit difference between the minimum and maximum intensity. Images were reconstructed with channel specific OTFs and aligned using the softWoRx 3D SIM image reconstruction software (Cytiva).

### Spermatid staging

To stage spermatids based on the shape of the nucleus, a line was drawn through the widest part of the nucleus. Spermatid cysts with nuclei wider than 3um were considered ‘Leaf,’ nuclei between 2 and 3um were considered ‘Late Leaf,’ and nuclei narrower than 2um were considered ‘Canoe.’

Asl labeling was used to stage spermatids based on the shape of the CA. Spermatids where Asl coated the length of the centriole were considered ‘centriolar,’ spermatids where Asl formed a collar over half the centriole were considered ‘collar,’ spermatids where Asl formed a ring were considered ‘ring,’ and spermatids where Asl was condensed to one side of the centriole were considered ‘remodeled.’

### Analysis of Spag4 embedding

Only spermatids where a Spag4 Cap was visible and where the centriole was in line with the nucleus were used. A line scan of the Spag4 signal was taken by drawing a 10 pixel wide by 3um long line centered on the middle of the Cap in Fiji. Background was eliminated by subtracting the average of the first five and last five intensity values from each value. Each intensity value was then normalized to the maximum intensity and the number of pixels which were more than 25% of the maximum was determined and then converted to microns.

### Analysis of centriole length

Only spermatids where centriole was parallel to the imaging plane were used. A line scan of the centriole (as labeled by PACT) was taken by drawing a 5 pixel wide by 3um long line centered on the middle of the centriole in Fiji. Background was eliminated by subtracting the average of the first five and lats five intensity values from each value. Each intensity value was then normalized to the maximum intensity and the number of pixels which were more than 50% of the maximum was calculated. This was then converted to microns to determine centriole length.

### Analysis of centriole lateralization

Spermatids were binned by whether they had only a Cap or a Cap and Shelf. A 5 pixel wide line was drawn across the width of the nucleus at the top of the Cap, perpendicular to the centriole in Fiji. In cases where there was a shelf, the beginning of the line was always set at the edge of the nucleus with the shelf. This line was moved down to the bottom of the nucleus and a line scan was taken through the centriole (PACT signal). The position of the peak of the centriole signal along this line maximum intensity was divided by the length of the line. If the centriole was perfectly centered in the nucleus, this would give a value of 0.5. If the centriole was positioned completely laterally, this would give a value of ∼1.

To determine the correlation between centriole lateralization and Shelf length, only spermatids with a visible Shelf were selected. The length of the Shelf was determined by drawing a line from the end of the Shelf to the edge of the centriole. The centriole lateralization was calculated as above and graphed relative to the Shelf length. Prism was used to determine the Pearson correlation coefficient.

### Analysis of PCL movement

Only spermatids where the PCL (as labeled by Sas-6) was in line with the centriole were used. A line scan of the Sas-6 signal was taken by drawing a 5 pixel wide by 2um long line through the sas6 signal along the centriole and centered between the proximal end of the centriole and the PCL in Fiji. Background was eliminated by subtracting the average of the first and last intensity values from each value. A sum of two Gaussians was performed in Prism and the peak to peak distance was calculated to determine the distance between the Sas-6 signal from the proximal end of the centriole to the Sas-6 from the PCL.

### Analysis of centriole insertion

Only spermatids where the centriole and nucleus could be identified were used. Centrioles that overlapped with the DAPI signal were considered ‘normal.’ Centrioles that were attached to and in line with the nucleus but did not overlap with the DAPI signal were considered ‘not inserted.’ Centrioles that were not inserted but were found at an irregular angle relative to the nucleus were considered ‘bent.’ Centrioles that were not in contact with the nucleus were considered ‘detached.’

### Analysis of Sas-6 fluorescence at the PCL

Only spermatids where the attachment of the centriole could be accurately assessed were used. A circular ROI was drawn around the Sas-6 signal at both the proximal end of the centriole and at the PCL in the Z-slice with the brightest signal. An ROI of the same size was taken from the adjacent background in the same slice and subtracted. Fluorescence of Sas-6 at the PCL was divided by the fluorescence at the proximal centriole end to generate a ratio. Spermatids were then binned by centriole insertion into one of the following categories: normal, not inserted, or detached.

### Statistics

For comparisons between two groups, unpaired *t* tests were used. Except where noted, one-way ANOVA with Tukey’s correction was used for comparisons between 3 or more groups. Sample sizes are denoted in the figure legends. All statistics were performed using GraphPad Prism 10.0.3. Scale bars represent the mean ± the standard deviation for each graph.

## REFERENCES

Anderson, M. A., Jodoin, J. N., Lee, E., Hales, K. G., Hays, T. S. & Lee, L. A. 2009. Asunder is a critical regulator of dynein-dynactin localization during Drosophila spermatogenesis. Mol Biol Cell, 20, 2709–21.

Anderson, W. A. 1967. Cytodifferentiation of spermatozoa in Drosophila melanogaster: the effect of elevated temperature on spermiogenesis. Mol Gen Genet, 99, 257–73.

Augiere, C., Lapart, J. A., Duteyrat, J. L., Cortier, E., Maire, C., Thomas, J. & Durand, B. 2019. salto/CG13164 is required for sperm head morphogenesis in Drosophila. Mol Biol Cell, 30, 636–645.

Baccetti, B., Burrini, A. G., Collodel, G., Magnano, A. R., Piomboni, P., Renieri, T. & Sensini, C. 1989. Morphogenesis of the decapitated and decaudated sperm defect in two brothers. Gamete Res, 23, 181–8.

Blachon, S., Cai, X., Roberts, K. A., Yang, K., Polyanovsky, A., Church, A. & Avidor-Reiss, T. 2009. A proximal centriole-like structure is present in Drosophila spermatids and can serve as a model to study centriole duplication. Genetics, 182, 133–44.

Blachon, S., Gopalakrishnan, J., Omori, Y., Polyanovsky, A., Church, A., Nicastro, D., Malicki, J. & Avidor-Reiss, T. 2008. Drosophila asterless and vertebrate Cep152 Are orthologs essential for centriole duplication. Genetics, 180, 2081–94.

Blachon, S., Khire, A. & Avidor-Reiss, T. 2014. The origin of the second centriole in the zygote of Drosophila melanogaster. Genetics, 197, 199–205.

Blom, E. & Birch-Andersen, A. 1970. Ultrastructure of the “decapitated sperm defect” in Guernsey bulls. J Reprod Fertil, 23, 67–72.

Chemes, H. E., Carizza, C., Scarinci, F., Brugo, S., Neuspiller, N. & Schwarsztein, L. 1987. Lack of a head in human spermatozoa from sterile patients: a syndrome associated with impaired fertilization. Fertil Steril, 47, 310–6.

Chemes, H. E., Puigdomenech, E. T., Carizza, C., Olmedo, S. B., Zanchetti, F. & Hermes, R. 1999. Acephalic spermatozoa and abnormal development of the head-neck attachment: a human syndrome of genetic origin. Hum Reprod, 14, 1811–8.

Demarco, R. S., Eikenes, A. H., Haglund, K. & Jones, D. L. 2014. Investigating spermatogenesis in Drosophila melanogaster. Methods, 68, 218–27.

Fabian, L. & Brill, J. A. 2012. Drosophila spermiogenesis: Big things come from little packages. Spermatogenesis, 2, 197–212.

Fuller, M. T. 1993. Spermatogenesis in Development of Drosophila, New York, Cold Spring Harbor Laboratory Press.

Galletta, B. J., Jacobs, K. C., Fagerstrom, C. J. & Rusan, N. M. 2016. Asterless is required for centriole length control and sperm development. J Cell Biol, 213, 435–50.

Galletta, B. J., Ortega, J. M., Smith, S. L., Fagerstrom, C. J., Fear, J. M., Mahadevaraju, S., Oliver, B. & Rusan, N. M. 2020. Sperm Head-Tail Linkage Requires Restriction of Pericentriolar Material to the Proximal Centriole End. Dev Cell, 53, 86–101 e7.

Ge, T., Yuan, L., Xu, L., Yang, F., Xu, W., Niu, C., Li, G., Zhou, H. & Zheng, Y. 2024. Coiled-coil domain containing 159 (CCDC159) is required for spermatid head and tail assembly in mice. Biol Reprod.

Jo, K. H., Jaiswal, A., Khanal, S., Fishman, E. L., Curry, A. N. & Avidor-Reiss, T. 2019. Poc1b and Sas-6 Function Together during the Atypical Centriole Formation in Drosophila melanogaster. Cells, 8.

Khanal, S., Leung, M. R., Royfman, A., Fishman, E. L., Saltzman, B., Bloomfield-Gadelha, H., Zeev-Ben-Mordehai, T. & Avidor-Reiss, T. 2021. A dynamic basal complex modulates mammalian sperm movement. Nat Commun, 12, 3808.

Khire, A., Jo, K. H., Kong, D., Akhshi, T., Blachon, S., Cekic, A. R., Hynek, S., Ha, A., Loncarek, J., Mennella, V. & Avidor-Reiss, T. 2016. Centriole Remodeling during Spermiogenesis in Drosophila. Curr Biol, 26, 3183–3189.

Khire, A., Vizuet, A. A., Davila, E. & Avidor-Reiss, T. 2015. Asterless Reduction during Spermiogenesis Is Regulated by Plk4 and Is Essential for Zygote Development in Drosophila. Curr Biol, 25, 2956–63.

Kierszenbaum, A. L., Rivkin, E. & Tres, L. L. 2011. Cytoskeletal track selection during cargo transport in spermatids is relevant to male fertility. Spermatogenesis, 1, 221–230.

Klebba, J. E., Buster, D. W., Nguyen, A. L., Swatkoski, S., Gucek, M., Rusan, N. M. & Rogers, G. C. 2013. Polo-like kinase 4 autodestructs by generating its Slimb-binding phosphodegron. Curr Biol, 23, 2255–2261.

Kracklauer, M. P., Wiora, H. M., Deery, W. J., Chen, X., Bolival, B., Jr., Romanowicz, D., Simonette, R. A., Fuller, M. T., Fischer, J. A. & Beckingham, K. M. 2010. The Drosophila Sun protein Spag4 cooperates with the coiled-coil protein Yuri Gagarin to maintain association of the basal body and spermatid nucleus. J Cell Sci, 123, 2763–72.

Li, M. G., Serr, M., Newman, E. A. & Hays, T. S. 2004. The Drosophila tctex-1 light chain is dispensable for essential cytoplasmic dynein functions but is required during spermatid differentiation. Mol Biol Cell, 15, 3005–14.

Martinez-Campos, M., Basto, R., Baker, J., Kernan, M. & Raff, J. W. 2004. The Drosophila pericentrin-like protein is essential for cilia/flagella function, but appears to be dispensable for mitosis. J Cell Biol, 165, 673–83.

Mcgee, M. D., Rillo, R., Anderson, A. S. & Starr, D. A. 2006. Unc-83 Is a Kash protein required for nuclear migration and is recruited to the outer nuclear membrane by a physical interaction with the Sun protein Unc-84. Mol Biol Cell, 17, 1790–801.

Padmakumar, V. C., Libotte, T., Lu, W., Zaim, H., Abraham, S., Noegel, A. A., Gotzmann, J., Foisner, R. & Karakesisoglou, I. 2005. The inner nuclear membrane protein Sun1 mediates the anchorage of Nesprin-2 to the nuclear envelope. J Cell Sci, 118, 3419–30.

Perotti, M. E., Giarola, A. & Gioria, M. 1981. Ultrastructural study of the decapitated sperm defect in an infertile man. J Reprod Fertil, 63, 543–9.

Platts, A. E., Dix, D. J., Chemes, H. E., Thompson, K. E., Goodrich, R., Rockett, J. C., Rawe, V. Y., Quintana, S., Diamond, M. P., Strader, L. F. & Krawetz, S. A. 2007. Success and failure in human spermatogenesis as revealed by teratozoospermic RNAs. Hum Mol Genet, 16, 763–73.

Riparbelli, M. G., Persico, V. & Callaini, G. 2020. The Microtubule Cytoskeleton during the Early Drosophila Spermiogenesis. Cells, 9.

Russell, L. D., Russell, J. A., Macgregor, G. R. & Meistrich, M. L. 1991. Linkage of manchette microtubules to the nuclear envelope and observations of the role of the manchette in nuclear shaping during spermiogenesis in rodents. Am J Anat, 192, 97–120.

Shoup, J. R. 1967. Spermiogenesis in wild type and in a male sterility mutant of Drosophila melanogaster. J Cell Biol, 32, 663–75.

Sitaram, P., Anderson, M. A., Jodoin, J. N., Lee, E. & Lee, L. A. 2012. Regulation of dynein localization and centrosome positioning by Lis-1 and asunder during Drosophila spermatogenesis. Development, 139, 2945–54.

Stanley, H. P., Bowman, J. T., Romrell, L. J., Reed, S. C. & Wilkinson, R. F. 1972. Fine structure of normal spermatid differentiation in Drosophila melanogaster. J Ultrastruct Res, 41, 433–66.

Stewart-Hutchinson, P. J., Hale, C. M., Wirtz, D. & Hodzic, D. 2008. Structural requirements for the assembly of Linc complexes and their function in cellular mechanical stiffness. Exp Cell Res, 314, 1892–905.

Tapia Contreras, C. & Hoyer-Fender, S. 2019. CCDC42 Localizes to Manchette, Htca and Tail and Interacts With ODF1 and ODF2 in the Formation of the Male Germ Cell Cytoskeleton. Front Cell Dev Biol, 7, 151.

Tates, A. D. 1971. Cytodifferentiation during spermatogenesis in Drosophila melanogaster: an electron microscope study’s Gravenhage.

Texada, M. J., Simonette, R. A., Johnson, C. B., Deery, W. J. & Beckingham, K. M. 2008. Yuri gagarin is required for actin, tubulin and basal body functions in Drosophila spermatogenesis. J Cell Sci, 121, 1926–36.

Tokuyasu, K. T. 1974a. Dynamics of spermiogenesis in Drosophila melanogaster. 3. Relation between axoneme and mitochondrial derivatives. Exp Cell Res, 84, 239–50.

Tokuyasu, K. T. 1974b. Dynamics of spermiogenesis in Drosophila melanogaster. Iv. Nuclear transformation. J Ultrastruct Res, 48, 284–303.

Tokuyasu, K. T. 1975a. Dynamics of spermiogenesis in Drosophila melanogaster. V. Head-tail alignment. J Ultrastruct Res, 50, 117–29.

Tokuyasu, K. T. 1975b. Dynamics of spermiogenesis in Drosophila melanogaster. Vi. Significance of “onion” nebenkern formation. J Ultrastruct Res, 53, 93–112.

Tokuyasu, K. T., Peacock, W. J. & Hardy, R. W. 1972a. Dynamics of spermiogenesis in Drosophila melanogaster. I. Individualization process. Z Zellforsch Mikrosk Anat, 124, 479–506.

Tokuyasu, K. T., Peacock, W. J. & Hardy, R. W. 1972b. Dynamics of spermiogenesis in Drosophila melanogaster. Ii. Coiling process. Z Zellforsch Mikrosk Anat, 127, 492–525.

Tokuyasu, K. T., Peacock, W. J. & Hardy, R. W. 1977. Dynamics of spermiogenesis in Drosophila melanogaster. Vii. Effects of segregation distorter (Sd) chromosome. J Ultrastruct Res, 58, 96–107.

Turner, K. A., Fishman, E. L., Asadullah, M., Ott, B., Dusza, P., Shah, T. A., Sindhwani, P., Nadiminty, N., Molinari, E., Patrizio, P., Saltzman, B. S. & Avidor-Reiss, T. 2021. Fluorescence-Based Ratiometric Analysis of Sperm Centrioles (Frac) Finds Patient Age and Sperm Morphology Are Associated With Centriole Quality. Front Cell Dev Biol, 9, 658891.

Wang, Y., Xiang, M. F., Zheng, N., Cao, Y. X. & Zhu, F. X. 2022. Genetic pathogenesis of acephalic spermatozoa syndrome: past, present, and future. Asian J Androl, 24, 231–237.

Zhang, Y., Liu, C., Wu, B., Li, L., Li, W. & Yuan, L. 2021. The missing linker between SUN5 and PMFBP1 in sperm head-tail coupling apparatus. Nat Commun, 12, 4926.

Zhu, F., Liu, C., Wang, F., Yang, X., Zhang, J., Wu, H., Zhang, Z., He, X., Zhang, Z., Zhou, P., Wei, Z., Shang, Y., Wang, L., Zhang, R., Ouyang, Y. C., Sun, Q. Y., Cao, Y. & Li, W. 2018. Mutations in PMFBP1 Cause Acephalic Spermatozoa Syndrome. Am J Hum Genet, 103, 188–199.

