## Supplemental Figures for "The Proximal Centriole-Like Structure Anchors the Centriole to the Sperm Nucleus"

Figure S1. The centriole and nucleus form a lateral attachment in mature spermatids

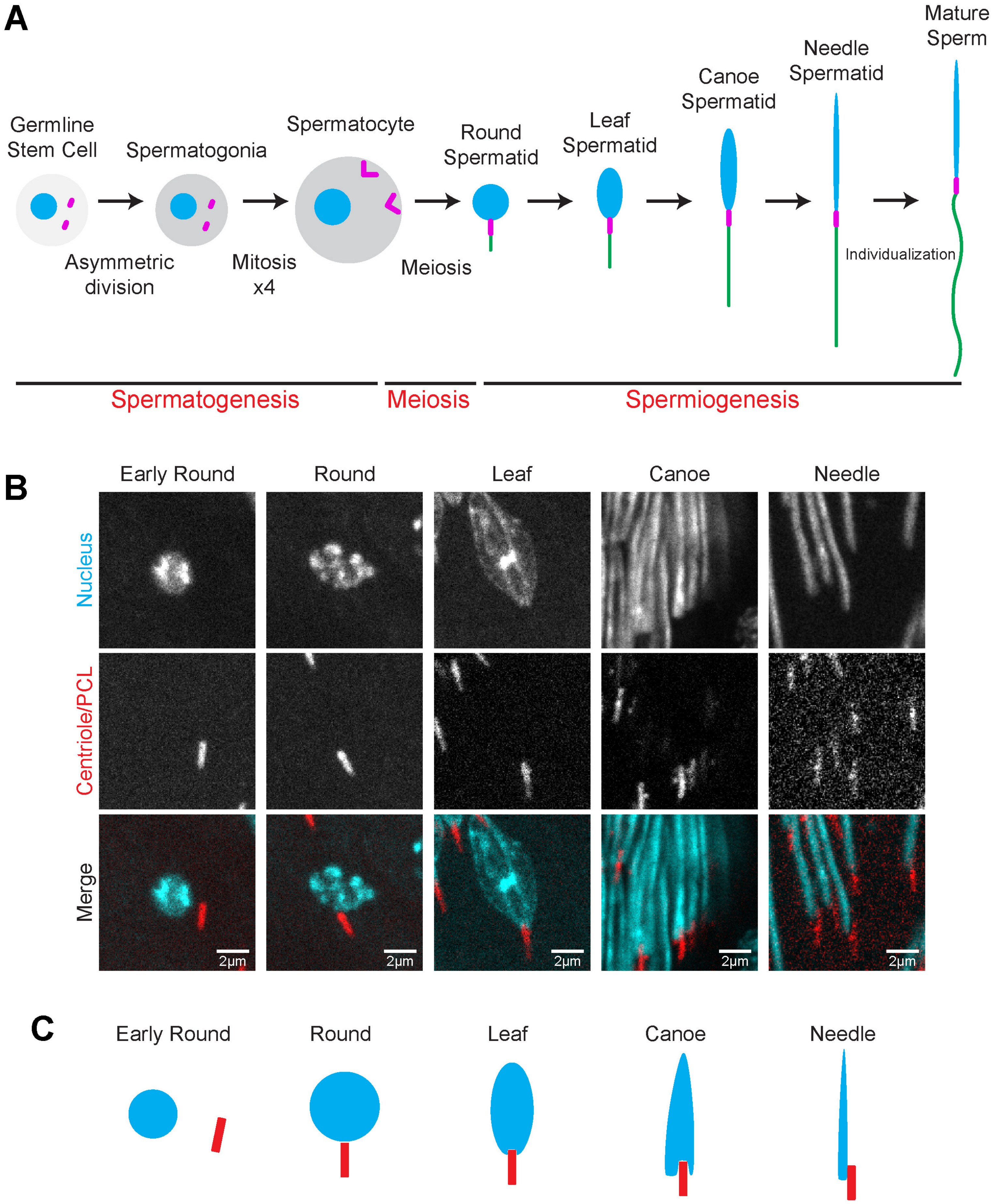

**Figure S1: The centriole and nucleus come together to form a lateral attachment in mature spermatids.**

**(A)** Cartoon depicting sperm development in *Drosophila*. In spermatogenesis, germline stem cells undergo an asymmetric division to generate gonioblasts (not depicted). These cells undergo 4 rounds of mitosis, when they are termed spermatogonia. Following these divisions, these cells become spermatocytes. Spermatocytes go through meios is to become spermatids. Spermatids undergo a series of dramatic morphological changes in which the nucleus (head) reshapes and the tail elongates in a process called spermiogenesis. **(B)** Representative images of spermatids at the indicated stages of spermiogenesis, beginning with Early Round spermatids through to Needle stage spermatids. Spermatids were labeled for the nucleus (DAPI, cyan) and the centriole/PCL (Ana1::tdTomato, red). Scale bar, 2µm. **(C)** Cartoons depicting the nucleus and centriole at each stage of spermiogenesis.

Figure S2. Yuri colocalizes with Spag4 at the Centriole Cap and Nuclear Shelf

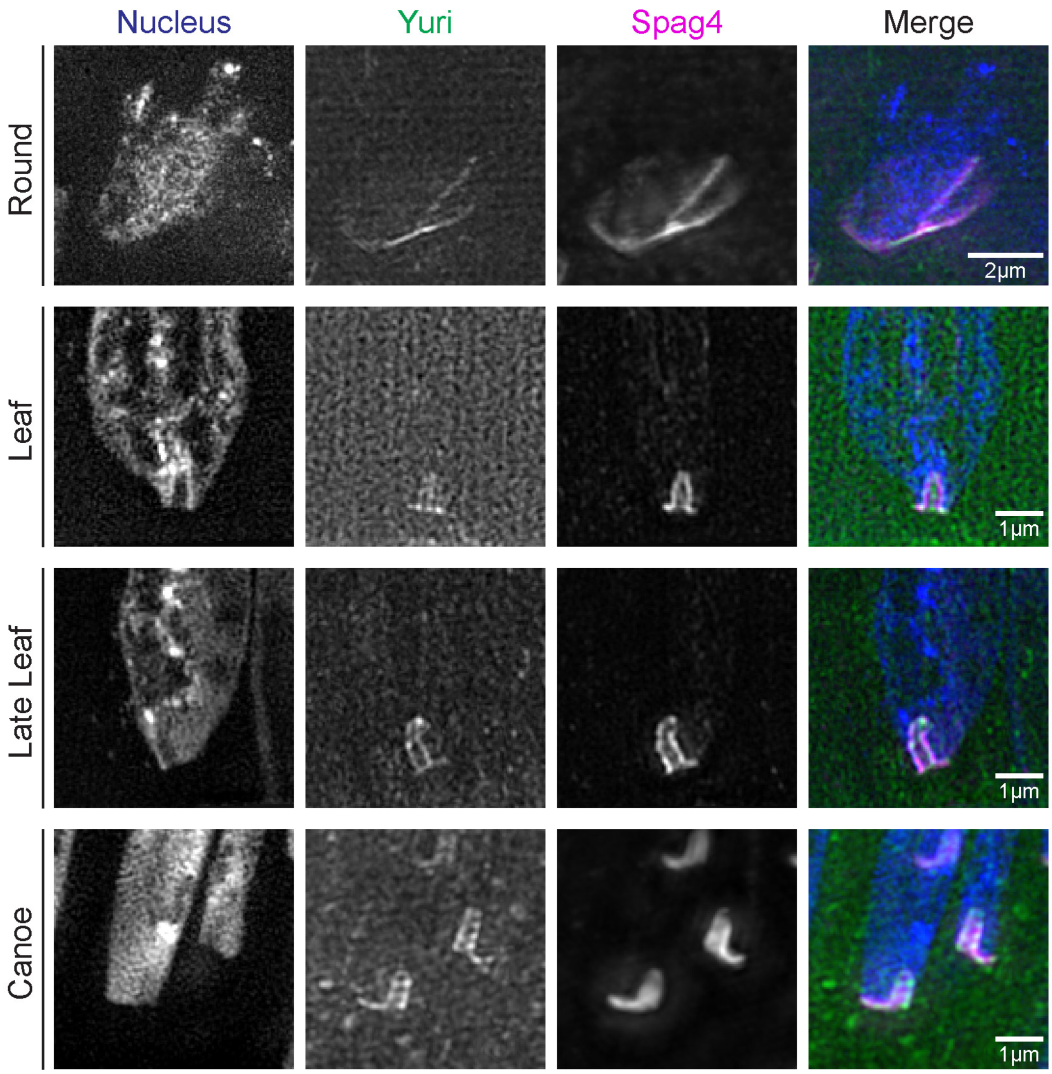

**Figure S2: Yuri colocalizes with Spag4 at the Centriole Cap and Nuclear Shelf**

Representative SIM images showing spermatids from wildtype testes during indicated developmental stages. Spermatids were labeled for the nucleus (DAPI, blue), Yuri (green), and Spag4 (Spag4::6myc, magenta). Scale bar, 2µm (Round), 1µm (Leaf, Late Leaf, and Canoe).

Figure S3. Poc1A and Poc1B localize to the PCL

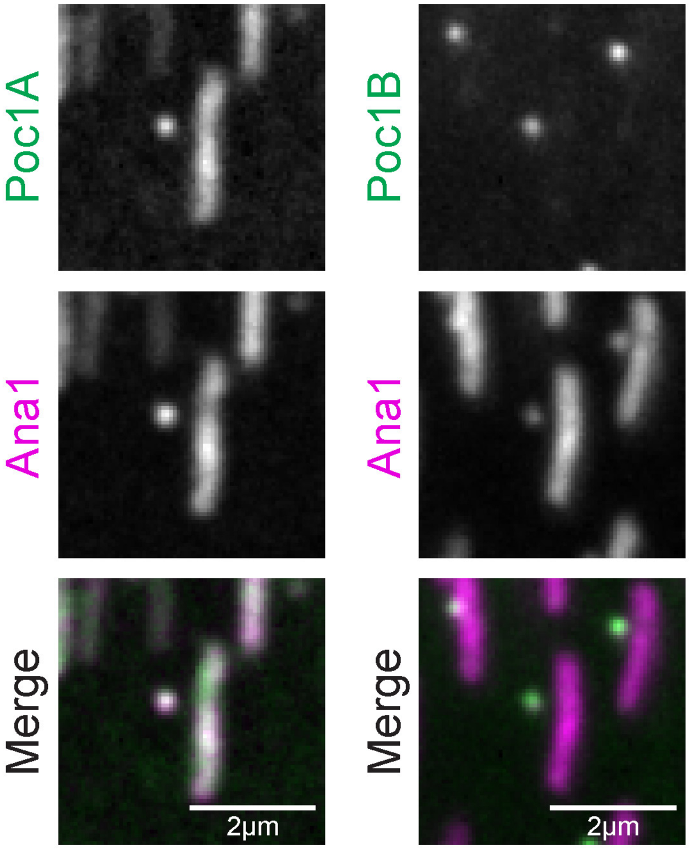

**Figure S3: Poc1A and Poc1B localize to the PCL.**

Representative images of centrioles from Canoe stage spermatids. Spermatids were labeled for the centriole/PCL (Ana1::tdTomato, magenta) and Poc1A (Poc1A::GFP, green, left) or Poc1B (Poc1B::GFP, green, right). Scale bar, 2µm.

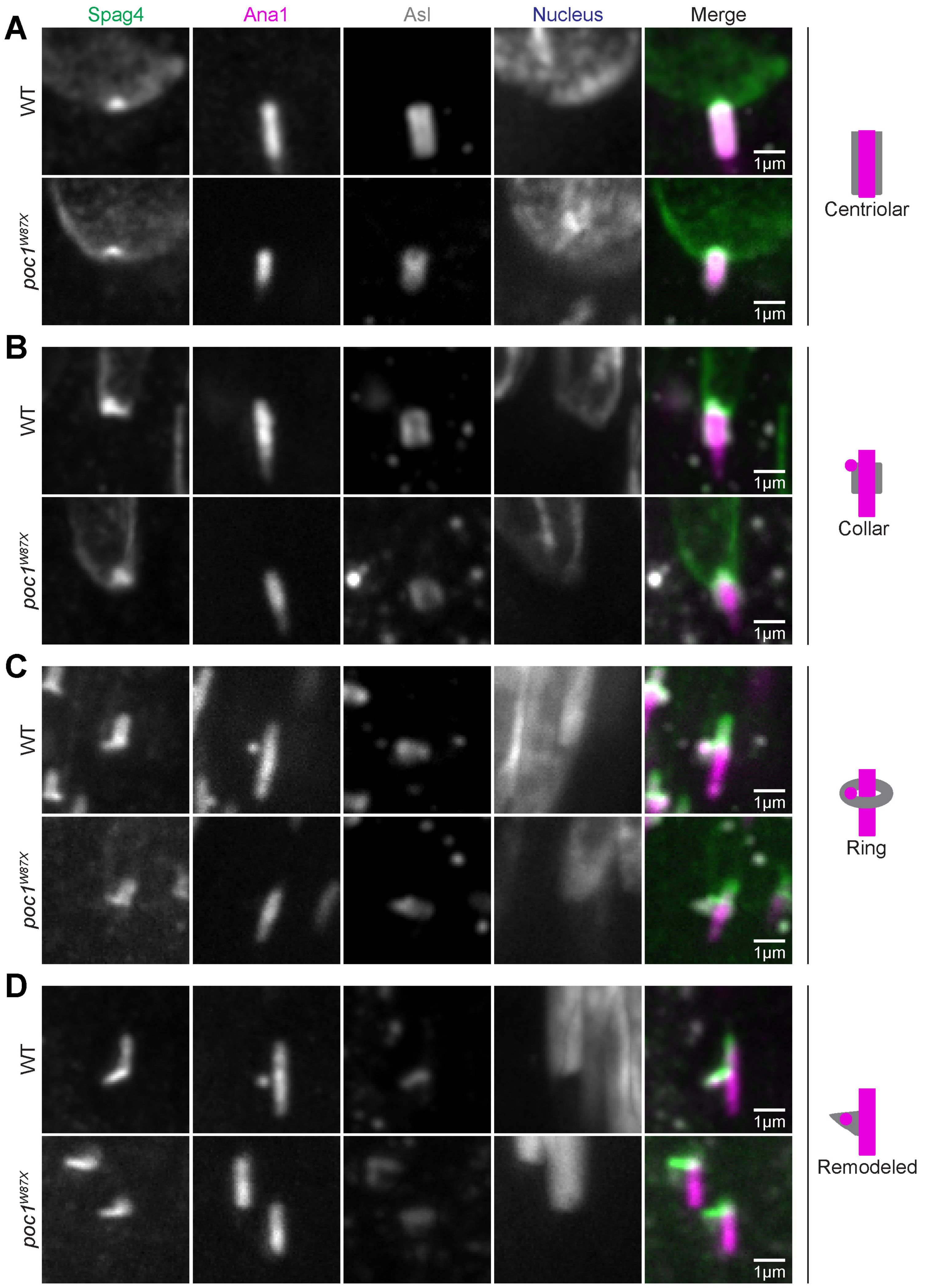


Figure S4. *poc1* mutants have shorter Centriole Caps following remodeling of the CA

**Figure S4: *poc1* mutants have shorter Centriole Caps following remodeling of the CA.**

**(A-D)** Individual channels for images of wildtype and *poc1* mutant spermatids shown in **Figure 4** at indicated stages of CA remodeling. Spermatids were labeled for the nucleus (DAPI, blue), Spag4 (Spag4::6myc, green), centriole (Ana1::tdTomato, magenta), and CA (Asl, gray). Scale bar, 1µm.

Figure S5. The PCL is required for proper centriole insertion

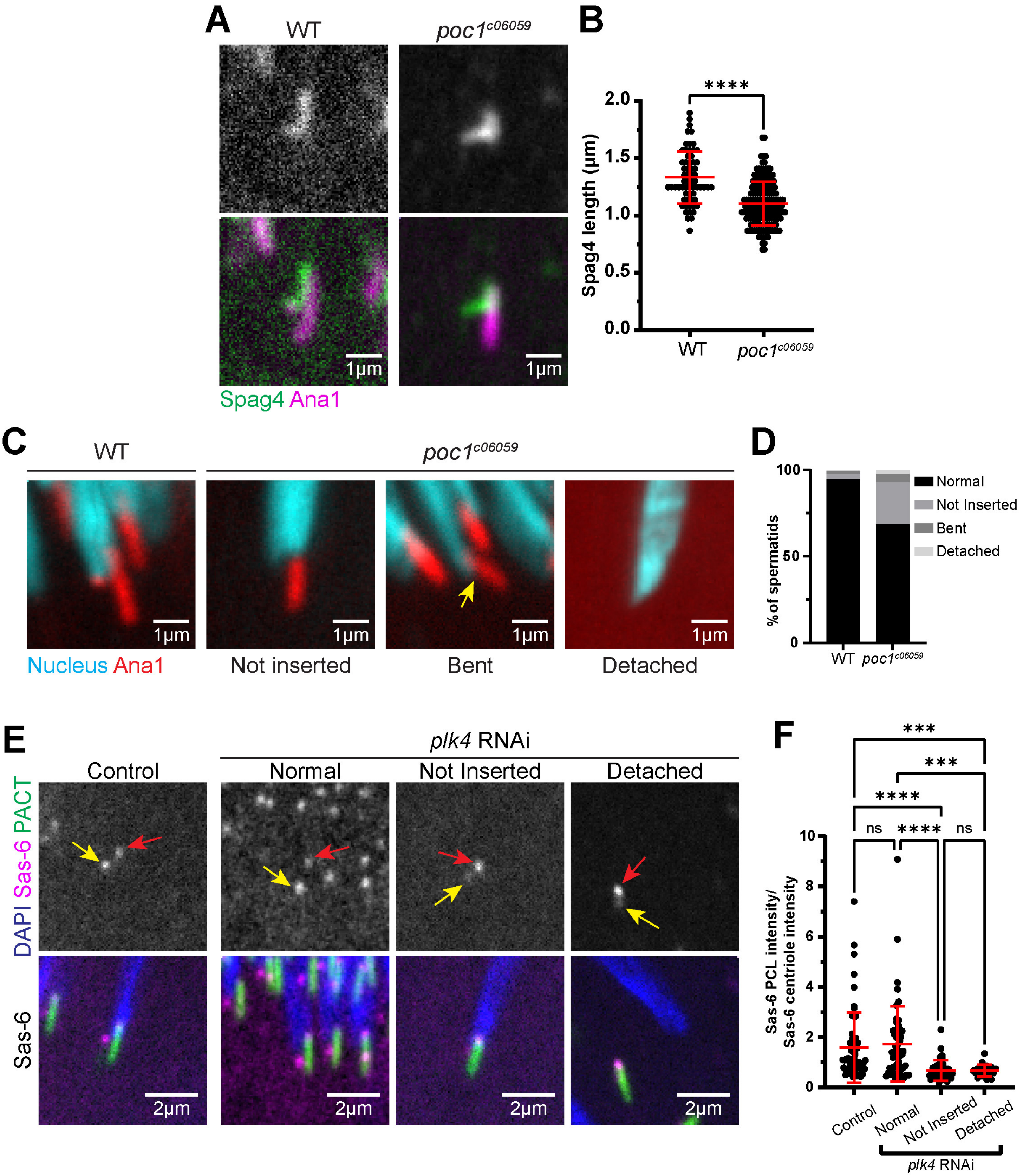

**Figure S5: The PCL is required for proper centriole insertion.**

**(A)** Representative images showing spermatids from wildtype (left) and *poc1*$^{c06059}$ mutant (right) testes at final stages of CA remodeling. Spermatids were labeled with Spag4 (Spag4::6myc, green) and the centriole/PCL (Ana1::tdTomato). Scale bar, 1µm. **(B)** Quantification of Centriole Cap length in wildtype and *poc1*$^{c06059}$ mutant spermatids during the remodeled (wildtype n=63; mutant n=163) stage of CA remodeling. (****) p≤0.0001. **(C)** Representative images showing a wildtype (left) spermatid with normal centriole insertion and a *poc1*$^{c06059}$ mutant (right) spermatid with a centriole that is not inserted. Spermatids were labeled with the nucleus (DAPI, cyan) and the centriole/PCL (Ana1::tdTomato, red). Yellow arrow denotes "bent" spermatid. Scale bar, 1µm. **(D)** Quantification of wildtype (n=128) and *poc1*$^{c06059}$ mutant (n=213) spermatids with various HTCA phenotypes. **(E)** Representative images showing Canoe stage spermatids from control (left) and *plk4* RNAi (right) with various HTCA phenotypes. Spermatids were labeled with the nucleus (DAPI, blue), centriole (PACT::GFP, green), and Sas-6 (TagRFP::Sas6, magenta). Yellow arrows denote Sas-6 signal at the PCL. Red arrows denote Sas-6 signal at the proximal end of the centriole. Scale bar, 2µm. **(F)** Quantification of Sas-6 signal at the PCL relative to Sas-6 signal at the proximal end of the centriole in control (n=59), *plk4* RNAi normal (n=52), *plk4* RNAi not inserted (n=38), and *plk4* RNAi detached (n=22) Canoe stage spermatids. ns=not significant, (***) p≤0.001, (****) p≤0.0001 by Kruskal-Wallis non-parametric test.
